# Model of neural induction in the ascidian embryo

**DOI:** 10.1101/2022.06.29.498205

**Authors:** Rossana Bettoni, Clare Hudson, Hitoyoshi Yasuo, Sophie de Buyl, Geneviève Dupont

## Abstract

How cell specification can be controlled in a reproducible manner is a fundamental question in development biology. In ascidians, a group of marine invertebrate chordates, geometry plays a key role in achieving this control. Here, we use mathematical modeling to demonstrate that geometry dictates the neural-epidermal cell fate choice in the 32-cell stage ascidian embryo by a two-step process involving first the modulation of ERK signaling and second, the expression of the neural marker gene, *Otx*. The model describes signal transduction by the ERK pathway that is stimulated by FGF and repressed by ephrin, and ERK-mediated control of *Otx* gene expression, which involves both an activator and an inhibitor of transcription. Considering the measured area of cell surface contacts with FGF- or ephrin-expressing cells as inputs, the solutions of the model reproduce the experimental observations about ERK activation and *Otx* expression in the different cells under normal and perturbed conditions. Sensitivity analyses and computations of Hill coefficients allow to quantify the robustness of the specification mechanism controlled by cell surface area and to identify the respective role played by each signaling input. Simulations also predict in which conditions the dual control of gene expression by an activator and an inhibitor that are both under the control of ERK can induce a robust ON/OFF control of neural fate induction.

**Author summary:** The development of a single cell zygote into a multicellular embryo occurs thanks to the combination of cell division and cell specification. The latter process corresponds to the progressive acquisition by the embryonic cells of their final physiological and functional characteristics, which rely on well-defined signaling-controlled genetic programs. The origin of the great robustness of cell specification remains poorly understood. Here, we address this question in the framework of the embryonic neural fate induction in ascidians, which are marine invertebrates. At the 32-cells stage, four cells identified by their precise location in the embryo adopt neural fate. On the basis of experimental observations, we develop a mathematical model that predicts that the choice between the neural or epidermal fate is controlled by the cell surface areas of the cells in contact with two antagonistic signals, FGF and ephrin. Our findings provide a computational confirmation of the major role played by the geometry of the embryo in controlling cell lineage acquisition during ascidian development.

## Introduction

Embryonic development is a reproducible process during which cells adopt different identities with a high spatiotemporal precision. This precision relies on the ability of cells to interpret signals from their environment and activate specific genetic programs. An accurate description of the interplay between signaling and transcriptional outputs is thus required to get a clear mechanistic knowledge of cell fate determination in embryonic development. In this context, mathematical models provide useful tools to develop a unifying framework to describe various types of experiments performed to investigate the relation between signaling and gene expression. Models can also answer questions that cannot be readily investigated experimentally (Sagner and Briscoe, 2017; Tosenberger et al., 2019).

Ascidians are marine invertebrate chordates belonging to the subphylum Tunicata, a sister group of vertebrates. Although the ascidian tadpole larvae exhibit the basic chordate body plan, these embryos are much simpler than their vertebrate cousins. Moreover, uniquely among chordate models, the embryogenesis of ascidians proceeds with an invariant cell division pattern, such that cellular configurations and cell cycle progression are quasi-invariant (Dumollard et al., 2013; Guignard et al., 2020). This allows precise identification of cells with known developmental outcomes. By combining experimental observations and computational modeling, Guignard et al. (2020) have shown that the cell contact areas between signal emitting and signal receiving cells could play a primary role in determining cell fates during cleavage and gastrula stage ascidian embryos. Geometric control of embryonic inductions contrasts with the role of morphogen gradients described in other systems, in which ligand concentrations determine cellular responses (Lander et al., 2002; Sagner and Briscoe, 2017).

In the present study, we focus on the neural-epidermal binary fate choice that takes place within the ectoderm field of the 32-cell stage ascidian embryo. This onset of neural induction is recognized by expression of the *Otx* gene. *Otx* is activated as an immediate-early response to the extracellular-signal-regulated kinase (ERK) pathway, downstream of fibroblast growth factor (FGF). ERK directly activates *Otx* expression via the ETS1/2 transcription factor (Bertrand et al., 2003). While the total 16 ectoderm cells are in direct contact with FGF-expressing vegetal cells (Fig. 1B) and are competent to respond to FGF, only four particular cells exhibit enough ERK activation levels to express *Otx* and thus adopt neural fate. The precision of this response depends on ephrin/Eph signaling taking place between ectoderm cells themselves (Ohta and Satou, 2013). We have recently shown that each ectoderm cell exhibits a level of ERK activation that correlates with its area of surface contact with FGF-expressing mesendoderm cells, while ephrin/Eph signals are important to dampen ERK activation across all ectoderm cells (Williaume et al., 2021). In contrast to the situation encountered *in Xenopus laevis* oocyte maturation for example (Ferrell and Machleder, 1998), during ascidian neural induction, ERK activation is not an all-or-none process but depends on the area of cell surface contact with FGF-expressing cells in a slightly supra-linear manner. The graded ERK activation response is converted into a bimodal transcription output of *Otx*, which is restricted to neural precursors. The transcription factor ERF2, also under the control of ERK activity, represses *Otx* expression in low-ERK cells. In consequence, the dual control exerted by ERK activity on the ETS1/2-mediated activation and ERF2-mediated repression of *Otx* expression appears to play the central role in generating ultrasensitivity in the ERK-*Otx* relationship, which allows neural induction to be an ON or OFF process (Williaume et al., 2021).

**Fig 1.**
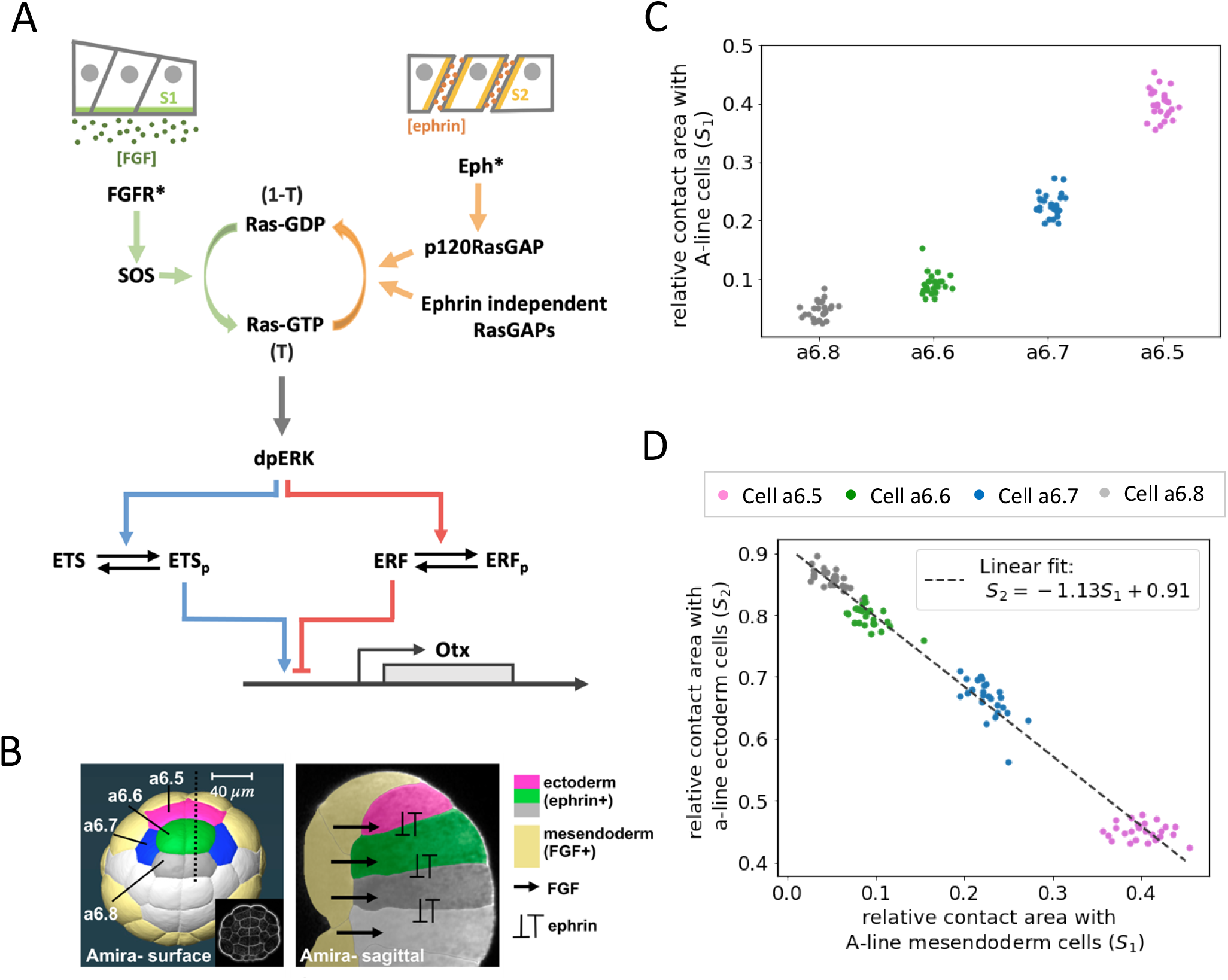
Model presentation. **(A)** a-line ectoderm cells of the ascidian embryo have different cell contact surfaces with FGF-expressing mesendoderm cells and with ephrin-expressing ectoderm cells. FGF binding activates the FGF receptor. Activated FGF receptos (FGFR*) then recruits the guanine nucleotide exchange factor SOS, which promotes the conversion of Ras-GDP to Ras-GTP. At the same time, the binding of the ephrin ligand, Efna.d, activates the Eph receptor. The active Eph receptor (Eph*) in turn recruits p120RasGAP, which promotes the conversion of Ras-GTP to Ras-GDP. Ras-GTP is also converted to Ras-GDP by ephrin-independent RasGAPs. Ras-GTP induces the activation (double phosphorylation: dp) of the extracellular signal-regulated kinase ERK via a kinase cascade including Raf and MEK. Phosphorylation of the ETS1/2 transcription factor by dpERK enhances the expression of the neural marker gene *Otx*. At the same time, dpERK phosphorylates the Ets repressor factor 2 (ERF2, indicated as ERF in the figure), which inhibits *Otx* expression. **(B)** Left: 32-cell stage ascidian embryo reconstructed with Amira software from a confocal z stack. a-line ectoderm cells are shown in different colors: cell a6.5 in magenta, cell a6.6 in green, cell a6.7 in blue, and cell a6.8 in gray. Mesendoderm cells are shown in yellow and b-line ectoderm cells in white. The dotted line is the line of sagittal section used for the right panel. Scale bar, 40 µm. Right: sagittal section highlighting cell surface contacts between ectoderm and mesendoderm cells and signals considered in the model. Figure from Willaume et al., 2021. **(C)** Relative area of a-line ectoderm cell surface in contact with A-line mesendoderm cells, S_1_, for the four cell types indicated in panel B. The relative contact area is computed as the surface contact with A-line/total cell surface. Each dot represents a single cell. **(D)** Relative area of a-line ectoderm cell surface in contact with ectoderm cells (S_2_) versus the relative contact area with A-line mesendoderm cells (S_1_). Fitting the experimental data with a linear function (shown as a black dashed line), we obtained the expression for the relation between S_1_ and S_2_: S_2_ = -1.13*S_1_+ 0.91 (R^2^= 0.9896). Figure from Willaume et al., 2021.

In our previous work (Williaume et al., 2021), we punctually used mathematical modeling to enhance the interpretation of the data and draw conclusions about some specific parts of the molecular mechanism underlying neural induction in the 32-cell stage ascidian embryo. In particular, minimal and non-parametrized modeling allowed the investigation of the role of cell surface contact *versus* ligand concentration in the ERK activation status, as well as the respective roles of the antagonistic enzymes SOS and p120RasGAP that control the regulatory status of Ras, the entry point for the ERK signaling cascade. A theoretical analysis of the dual control of *Otx* expression by an activator and a repressor, both under the control of ERK activity, was also performed to investigate the possible effect of a repressor on the shape of the ERK-*Otx* relation. Here, we combine the different mathematical modules developed previously to provide an enhanced, full mathematical description of FGF and ephrin-signaling control of *Otx* expression during early neural fate induction in the 32-cell stage ascidian embryo. The model, which consists of a set of ordinary differential equations, is solved at steady state. Results indicate that a unique set of parameters can adequately account for experimental observations related to neural induction when cells of the ascidian embryo are supposed to differ only by their cell surface contacts with FGF- and ephrin-expressing cells. The same set of parameter values can account for observations performed on pharmacologically or experimentally perturbed embryos. The robustness of the model is corroborated by sensitivity analysis. Finally, the model is used to make theoretical predictions by exploring relations that have not yet been obtained experimentally, for example the dependence of *Otx* expression on the cell surface area in contact with the signaling molecules in the absence of the ERF2 repressor.

## Results

The model takes into account that each cell perceives a level of FGF signaling that is proportional to the area of cell contact with FGF-expressing mesendoderm cells and a level of ephrin signaling that is proportional to the area of cell contact with ephrin-expressing ectoderm cells, as schematized in Fig. 1A. These surfaces have been quantified (Fig. 1B-1D) and are used to determine the number of receptors in contact with the signals. Ligand-bound FGF receptors (FGFR) and ephrin-bound Eph receptors stimulate the activity of SOS and p120RasGAP, respectively. SOS transforms Ras-GDP into Ras-GTP. Ras-GTP is converted back into Ras-GDP by p120RasGAP and by a cell surface contact-independent GAP activity. Ras-GTP activates the ERK signaling cascade. The relation between ERK activity and the level of Ras-GTP is described by a single function, which represents the full Ras-Raf-MEK-ERK pathway. Active ERK in turn phosphorylates the two transcription factors, ETS1/2 and ERF2. Phosphorylated ETS1/2 promotes the expression of the neural immediate-early gene *Otx*, while unphosphorylated ERF2 represses this expression. The model consists in 5 ordinary differential and 13 algebraic equations describing the abovementioned phenomena using Michaelis-Menten and Hill type kinetic expressions. A full description of the model is provided in the Modeling section.

In the 32-cell stage ascidian embryo, neural induction takes place in the ectoderm lineages (anterior (a) and posterior (b) lines) and is dependent on cell-cell contact with FGF-expressing mesendoderm cells (anterior (A) and posterior (B) lines). We focus on the four pairs of a-line cells, namely a6.5, a6.6, a6.7, and a6.8 (see Fig. 1B; Williaume et al., 2021). The four cells display different relative areas of cell surface contact with A-line mesendoderm cells that express FGF, with the neural precursor, a6.5, having the largest area of cell surface contact (Fig. 1C). Areas of cell surface contact with FGF-expressing cells and ephrin-expressing cells are inversely correlated, such that cells having the largest cell surface contact with FGF-expressing cells have the lowest cell surface contact with ephrin-expressing ectoderm cells (Fig. 1D). In the following, we model ERK activation and *Otx* expression in these four cells.

### Regulation of ERK activity by fractions of cell surfaces exposed to antagonistic signals

Experimental evidence indicates that, in the 32-cell stage ascidian embryo, the level of ERK activition in ectoderm cells is controlled by the extent of cell surface in contact with FGF-expressing mesendoderm cells and with ephrin-expressing ectoderm cells (Williaume et al., 2021). Because these two contact areas are correlated, the area of cell surface contact with FGF-expressing cells (designated S_1_) can also be used to define the area of cell surface contact with ephrin-expressing cells (designated S_2_). Thus, in terms of modeling, the only parameter that is cell-type-specific is S_1_. The steady states solutions of eqs. (1-13) should correspond to the levels of ERK activation measured in the different cell types as anti-dpERK immunofluorescence (IF) signals in their nucleus. Importantly, extracellular FGF and ephrin concentrations are assumed to be the same for all cell types. As shown in Fig. 2A, good agreement between observed and modeled ERK activation levels can be obtained for the four cell types when considering the values of parameters listed in Table 1 and assuming a linear relationship between computed normalized ERK activity (Erk*) and dpERK-IF signal measured experimentally (Eq 14).

**TABLE 1.** Default values of parameters. These values were obtained by manual fitting to get best agreement with experimental observations.

| Parameter | Definition | Value |
| --- | --- | --- |
| $A$ | Maximal level of dpERK fluorescence | Variable |
| $B$ | Basal level of dpERK fluorescence | Variable |
| $C$ | Maximal number of smFISH Otx spots | Variable |
| $D$ | Basal smFISH Otx spots count | Variable |
| $[\text{ephrin}]$ | Concentration of ephrin | 5 |
| $[\text{FGF}]$ | Concentration of FGF | 5 |
| $k$ | Degradation rate for Otx | 0.2 |
| $K_a$ | Half saturation constant for the activator ETS1/2 | 0.1 |
| $K_b$ | linear reaction rate of conversion from Ras-GTP into Ras-GDP due to contact surface independent RasGAP activity | 0.2 |
| $K_d$ | Binding constant of FGF to its receptor FGFR | 25 |
| $K_e$ | Binding constant of ephrin to its receptor Eph | 50 |
| $K_{erk}$ | Fraction of Ras-GTP leading to half maximal dpErk activity | 0.5 |
| $K_i$ | Dissociation constant for the inhibitor ERF2 | 0.1 |
| $k_{MM1}$ | Rate constant for the phosphorylation reaction of the inhibitor ERF2 | 12 |
| $K_{MM1}$ | Michaelis-Menten constant for the phosphorylation of $I$ (ERF2) | 0.05 |
| $K_{MM2}$ | Michaelis-Menten constant for the dephosphorylation of $I_p$ (phosphorylated ERF) | 0.05 |
| $k_{MM3}$ | Rate constant for the phosphorylation reaction of the activator ETS1/2 | 12 |
| $K_{MM3}$ | Michaelis-Menten constant for the phosphorylation of $A$ (ETS1/2) | 0.05 |
| $K_{MM4}$ | Michaelis-Menten constant for the dephosphorylation of $A_p$ (phosphorylated ERF2) | 0.05 |
| $K_{rg}$ | Number of ephrin-bound Eph receptors leading to half of the maximal p120RasGAP activity | 1200 |
| $K_s$ | Number of FGF-bound FGF receptors leading to half of the maximal SOS activity | 1200 |
| $K_1$ | Normalized half-saturation constant of SOS for its substrate Ras-GDP | 0.5 |
| $K_2$ | Normalized half-saturation constant of p120RasGAP for its substrate Ras-GTP | 0.2 |
| $n$ | Hill coefficient for the ERK pathway | 2 |
| $Q_T$ | Total number of ephrin receptors present on the cell membrane | 2000 |
| $R_T$ | Total number of FGF receptors present on the cell membrane | 2000 |
| $S_1$ | Fraction of cell surface contact with FGF | Variable |
| $S_2$ | Fraction of cell surface contact with ephrin | Variable |
| $v_b$ | Basal *Otx* expression rate | 0.001 |
| $v_{MM2}$ | Maximal rate at which $I_p$ (phosphorylated ERF2) is dephosphorylated | 1 |
| $v_{MM4}$ | Maximal rate at which $A_p$ (phosphorylated ETS1/2) is dephosphorylated | 1 |
| $v_o$ | $v_b + v_o$ is the maximal *Otx* expression rate | 1 |
| $V_s$ | Maximal rate of SOS activation | 1 |
| $V_{rg}$ | Maximal rate of p120RasGAP activation | 0.4 |

**Fig 2.**
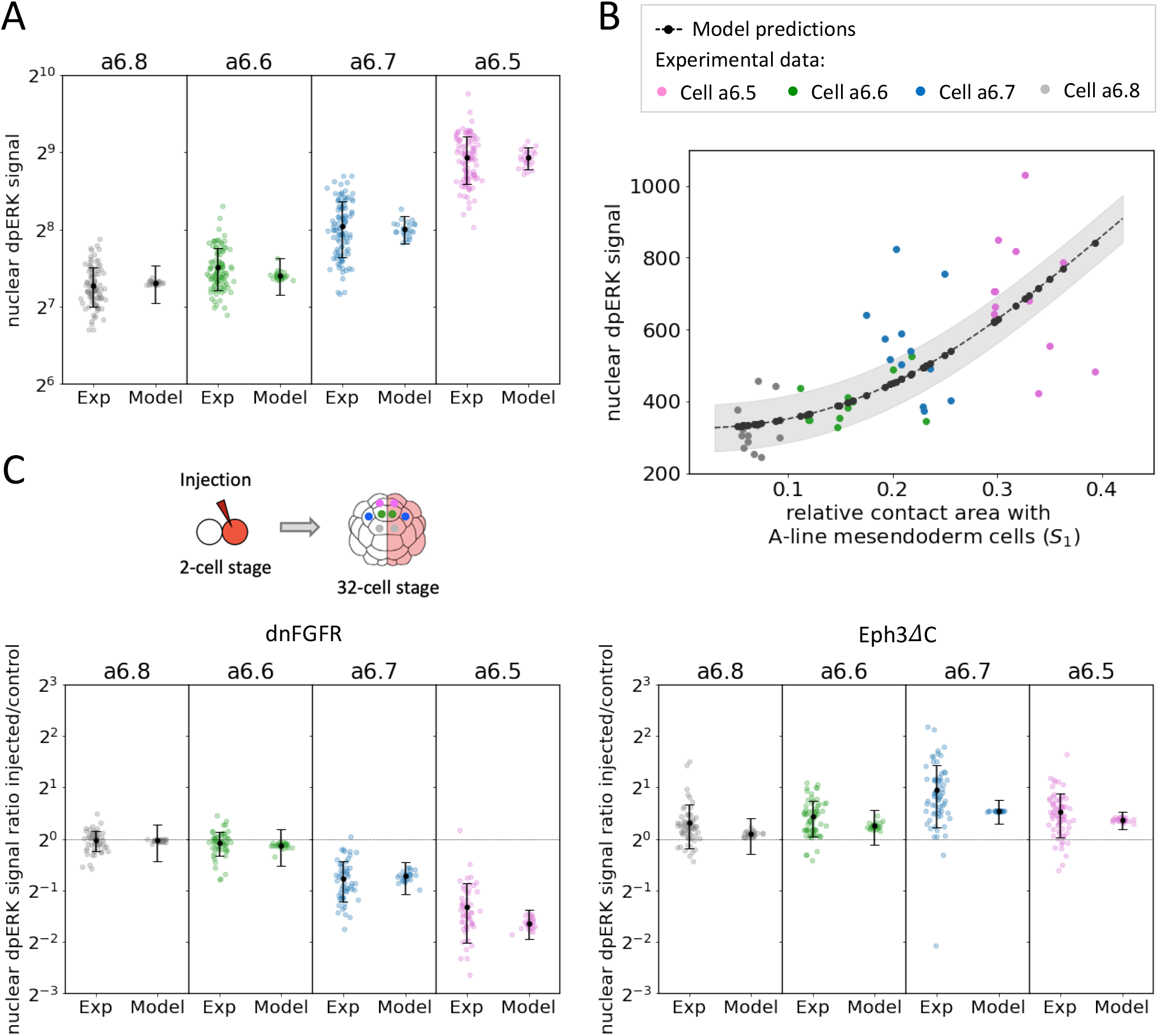
Control of ERK activation by cell contact surfaces. **(A)** Nuclear dpERK signals in the a6.5, a6.6, a6.7 and a6.8 cell types as measured in IF experiments (left) and computed with the model (right, *Erk*^*f*^ values). Each point represents a single cell and modeling results are computed using the measured values of S_1_. Means and standard deviations are shown in black. A=1850 and B= 155.11 in Eq 14. **(B)** Nuclear dpERK IF signals in individual a-line cells are shown as a function of the relative area of cell surface contact with A-line cells in experiments and in the model. Experimental data are shown in grey (a6.8 cell type), green (a6.6 cell type), blue (a6.7 cell type), magenta (a6.5 cell type), predictions of the model in black. Hill coefficient obtained by fitting the model predictions with a Hill function: 2.41. A=3000 and B= 324.52 in Eq 14. The area shaded in grey represent the uncertainty on the model prediction. **(C)** Injected/control ratios of nuclear dpERK signal in dnFGFR half-injected embryos (on the left) and in Eph3ΔC half-injected embryos (on the right). Left and right columns show experimental dpERK and computed *Erk*^*f*^ ratios, respectively. Injection of dnFGFR and Eph3ΔC were modeled by considering R_T_=100 and Q_T_=10, respectively. A= 1850 and B=155.11 in Eq 14.

The computed Erk* activities are equal to 0.0021 ± 0.0013, 0.0078 ± 0.0038, 0.0551 ± 0.0096 and 0.1787 ± 0.0201 in the a6.8, a6.6, a6.7 and a6.5 cell types, respectively (Fig. S1A). Each value was computed as the average between the steady state solutions obtained for the 25 values of the area of surface contact measures experimentally. Computed values reproduce the observed moderately nonlinear relation between ERK activation and cell surface in contact with FGF-expressing mesendoderm cells. The nonlinear relationship between S_1_ and ERK activity can also be compared with experiments where cell surfaces and dpERK were both measured in individual cell level (Fig. 2B, n_H_=2.41). The model predicts a modest level of ERK activation, defined between 0 and 1, in the maximally activated a6.5 cell, ∼0.18 (Fig. S1D). This value stems from the assumed relatively low concentration of extracellular FGF (K_D_/5) and, to a lesser extent, from the presence of ephrin (see below).

Much of the uncertainty when comparing Erk* to measured dpERK IF signals stems from the basal level of fluorescence in experimental measurements. Experimentally determined values are scattered around computed ones (Fig. 2). It remains unknown how much of the variation in measured ERK levels within each cell type are true to the system or are inherent to the experimental technique. The larger dispersion of experimental than theoretical values may reveal the existence of cell-to-cell heterogeneities that are not considered in the model or reveal intrinsic noise due to molecular fluctuations (see discussion).

FGF and ephrin control the activation of the ERK pathway during early neural induction and could thus both govern the emergence of the neural cell fate. Because of the inverse relationship between the cell surface contact with FGF-expressing mesendoderm cells and that with ephrin-expressing ectoderm cells (Fig. 1D), the a6.5>a6.7>a6.6>a6.8 profile of ERK activation levels in the 32-cell stage *Ciona* embryo could rely either on an increasing gradient of activation of SOS or on a decreasing gradient of inactivation of p120RasGAP. The respective contributions of these opposite influences were investigated experimentally by several targeted manipulations of the two pathways and by modeling (Williaume et al., 2021). It was shown that the profile of ERK activation can be controlled by the gradual activation of FGF receptors, even in the absence of ephrin signals. Ephrin signaling is important to reduce the levels of ERK activation across all ectoderm cells. Modeling predicted a greater influence of SOS compared to p120RasGAP on ERK activation levels. In the following, we validated the model and the parameter values by simulating the various experiments conducted in Williaume et al (2021).

The pharmacological inhibitor of ephrin receptors, NVP-BHG712 (henceforth NVP), has been shown to inhibit Eph signals in ascidian embryos (Fiuza et al, 2020; Williaume et al., 2021). When the model equations were solved with a minimal value of ephrin (0.001) to simulate the presence of the inhibitor, the a6.5>a6.7>a6.6>a6.8 profile of Erk* activation in the four cell types remained unchanged (Fig. S1C), as observed experimentally. Moreover, the model still exhibits a positive correlation between the area of cell surface contact with FGF-expressing cells and ERK activation levels, although the dependence is smoother and less nonlinear (Fig. S1B, n_H_=1.96). The effect of altering the SOS pathway was investigated experimentally by injecting a dominant negative form of the FGF receptor (dnFGFR) into one of the cells of the two-cell stage embryo. In these conditions, dnFGFR blocks FGF signals in one half of the embryo at the 32-cell stage, while the non-injected side represents the control situation. The effect of dnFGFR was simulated by decreasing the number of FGF receptors (*R*_*T*_) by 20 (i.e. *R*_*T*_ = 100). Observed ratios of dpERK-IF signals between injected and control sides are shown in Fig. 2C. Computations predict that the decrease in the number of FGF receptors leads to a marked decrease in the ERK activity in all cell types: Erk* now ranges from ∼5 10^−6^ in a6.8 cells to ∼5 10^−4^ in a6.5 cells. Thus, the ERK pathway is practically totally inactive in all cell types when FGFR is inhibited. In a symmetrical manner, the model reproduces the effect of the injection of a dominant negative form of the Eph3 receptor (Eph3ΔC) (Fig. 2C). Best agreement with experimental data is found when dividing the number of ephrin/Eph receptors (Q_T_) by 200 (i.e. Q_T_ = 10).

In summary, simple phenomenological equations describing the regulation of the ERK signaling pathway by FGF receptor-controlled SOS activation and ephrin/Eph-controlled RasGAP inactivation can quantitatively account for experimental observations in the 32-cell stage ascidian embryo. Steady-state solutions of the model confirm that gradual ERK activation levels are controlled by the different cell surface contacts to FGF and ephrin-expressing cells. The relation between these surfaces and ERK activation levels is moderately nonlinear. The model predicts that the control exerted by the FGF-receptor pathway is stronger than that exerted by ephrin/Eph. This assumption cannot be directly tested experimentally. Indeed, while the effect of FGF on ERK activation can be tested by inhibiting the ephrin pathway, the effect of ephrin in the absence of FGF cannot be evaluated because the level of ERK activity would be undetectable in all cells. The model also predicts that the ERK pathway is far from being maximal at the 32-cell stage of the ascidian embryo, even in the most-strongly activated a6.5 cell type.

### Dual control of the expression of the Otx gene by ERK

The *Otx* gene is a direct target of ERK signals during the initial step of ascidian neural induction (Bertrand et al, 2003; Hudson & Lemaire, 2003). Our previous results (Williaume et al., 2021), modeled in the preceding section, have shown that ERK activation in the 32-cell ascidian embryo exhibits a relatively smooth profile in response to contact-dependent exposure to FGF signals (Fig. 2A,B). In contrast, the transcription of the immediate-early gene *Otx* is bimodal, with a ON response that is restricted to neural precursors. We hypothesized that the ON/OFF response is, at least in part, due to the dual control of *Otx* expression (Williaume et al., 2021): ETS1/2 phosphorylation by ERK is required for its transcriptional activator activity (Weaver et al., 2022), while phosphorylation of the ERF2 repressor by dpERK promotes its nuclear export, thus preventing its binding to DNA (Le Gallic et al., 1999; 2004) (Fig. 1A).

To further validate this hypothesis, we have extended the model of ERK activation studied in the previous section to include the relation between ERK activition, phosphorylated ETS1/2, unphosphorylated ERF2 and *Otx* expression. We first seek a single set of parameter values accounting for the observed levels of *Otx* expression in the four cell types. The existence of such a set would corroborate the assumption that the emergence of the neural cell fate in *Ciona* is controlled by the contact areas with FGF and ephrin-expressing cells. Steady state levels of *Otx* smFISH spots computed with the model defined by eqs (1-18), using the parameter values listed in Table 1, are shown in Fig. 3A, together with experimental data for comparison (see also Fig. S2A for the O values before the linear transformation into *Otx* smFISH spots). When simulating the inhibition of FGF signaling (R_T_ = 100 instead of 2000), O values drop to nearly zero in all cell types, which corresponds to the absence of *Otx* expression observed in dnFGFR-injected half embryos (Fig. 3B). In the model as in the experiments, *Otx* expression is enhanced by inhibition of Eph signaling. This is noticed in the simulations of invalidation of the expression of ephrin/Eph receptors (Q_T_ = 10 instead of 2000) or of p120RasGAP enzymes (V_rg_ = 0.01 instead of 0.4), or by simulating the inhibition of the ephrin/Eph receptors by NVP ([ephrin]=0.001 instead of 5) (see Fig. 3B and 3C). In all cases, these treatments reproduce the observed ectopic activation of the a6.7 cells. This suggests that the moderate increase in ERK activition that results from the absence of Eph signaling (Fig. 2C right and Fig. S1C) has a drastic effect on *Otx* expression in this cell type. In Eph-inhibited embryos, if ERK signaling is reduced by another means (low dose of the U0126 inhibitor that is simulated by increasing K_erk_), an *Otx* output similar to the wild-type pattern is recovered (Fig. 3C, right column).

**Fig 3.**
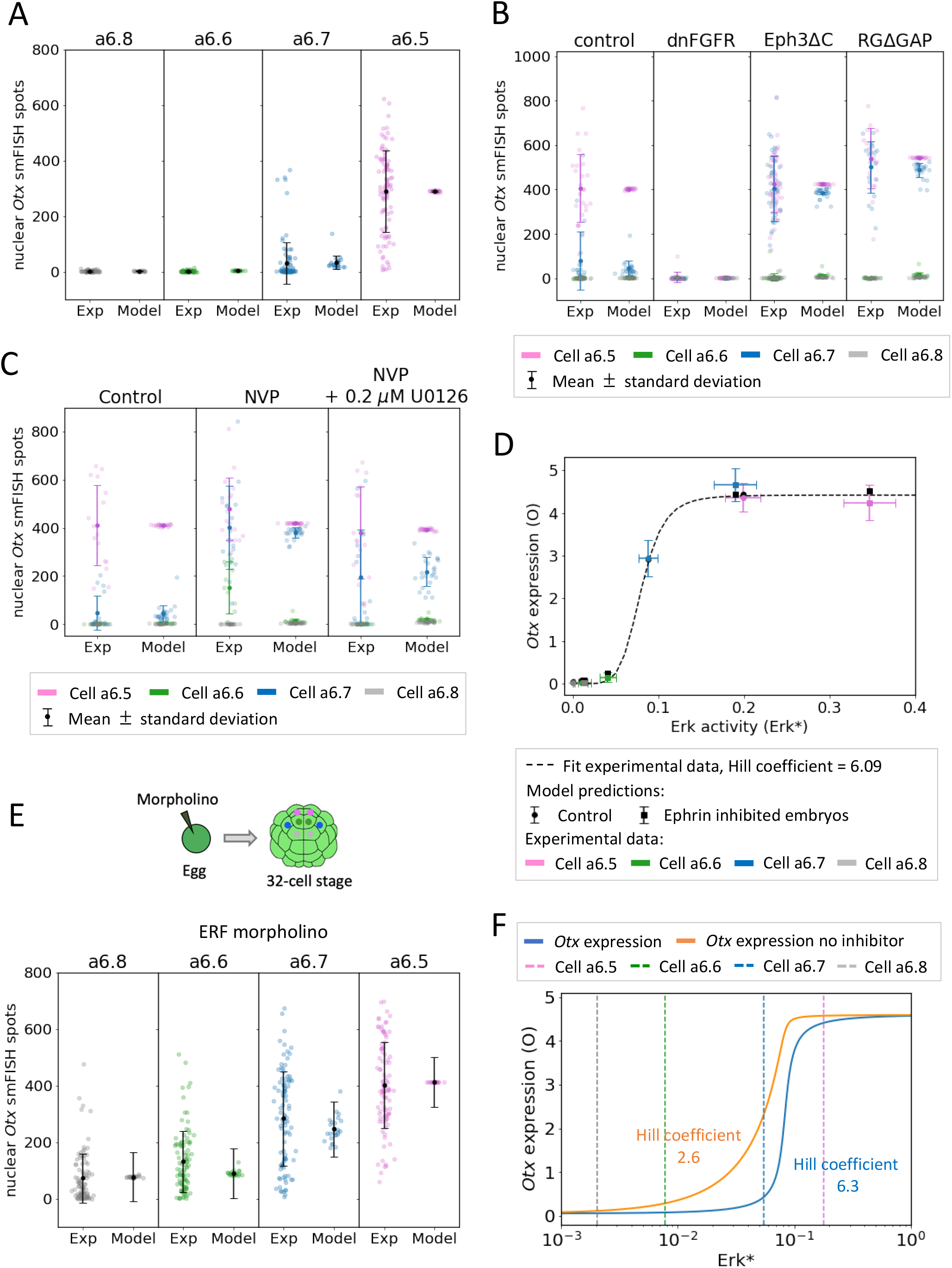
Control of *Otx* expression by ERK. **(A)** Levels of *Otx* expression in the a6.5, a6.6, a6.7 and a6.8 cell types as measured by single molecule fluorescence *in situ* hybridization (smFISH, left) and computed with the model (right, *Otx*_*smFISH*_). Each point represents a single cell and modeling results are computed using the measured values of S_1_. Means and standard deviations are shown in black. C=66 and D=2.75 in Eq 18. **(B)** *Otx* expression in control, dnFGFR, Eph3ΔC and RGΔGAP injected embryo halves. Left and right columns show experimental *Otx* smFISH spots and computed *Otx*_*smFISH*_, respectively. Injection of dnFGFR, Eph3ΔC and RGΔGAP were modeled by considering R_tot_=100, Q_tot_=10, and V_rg_=0.01, respectively. In Eq 18, C=92 for the control embryos and for dnFGFR injected embryos, C=95 for Eph3ΔC injected embryos and C=122 for RGΔGAP injected embryos. D=1.5 for the control and dnFGFR injected embryos, D=2.71 for Eph3ΔC and D=1.61 for RGΔGAP injected embryos. **(C)** Effect of the ephrin/Eph inhibitor NVP on *Otx* expression in the four cell types. On the left, control; in the middle, NVP-treated embryos; on the right, embryos treated with NVP and with the ERK inhibitor U0126 0.2 *µ*M. In the three cases, experimental *Otx* smFISH spots are shown on the left and computed *Otx*_*smFISH*_ values on the right. Results for the four cell types are shown in the same column with different colors. Each dot represents a single cell. NVP and moderate U0126 treatment are simulated by considering [ephrin]= 0.001 and K_erk_ =1, respectively. In Eq 18, C = 94, D=1.2. **(D)** *Otx* expression as a function of ERK activity (Erk*) in the four cell types (colored points and squares) and computed with the model (*Otx*_*smFISH*_, black points and squares). In the two cases, dots indicate the control while squares indicate the NVP-treated embryos. The experimental data were fitted by a Hill function, best-fit Hill coefficient = 6.09. For the experimental points, the value of Erk* corresponding to each cell was obtained by inversion of Eq 14, with A=3200 and B=0. The experimental value of O corresponding to each cell was obtained by inversion of Eq 18, with C=120 and D=0. Modelled Otx outputs were computed using the experimentally measured Erk* estimates as inputs. **(E)** Levels of *Otx* expression in the a6.5, a6.6, a6.7 and a6.8 cell types when unfertilized eggs have been injected with the ERF2-morpholino to prevent translation of ERF2. Injection of this morpholino is modeled by considering I as a constant equal to 0.01 in Eq 15. C=75 and D=73.4 in Eq 18. Modeling results for control embryos are the same as in panel (A). **(F)** *Otx* expression (O) as function of Erk*, computed with the model considering the presence (blue line, Hill coefficient = 6.3) or the absence (orange line, Hill coefficient= 2.6) of I (inhibitor). The Hill coefficients were computed using relation (22). Dashed vertical lines represents the mean values of Erk* for each cell type.

In fact, a highly nonlinear relationship between *Otx* expression and ERK activity is observed in the experiments and in the model. In Fig. 3D, values of Erk* and O corresponding to the experiments are inferred by inversion of eq. (14) and (18), respectively. Experimental values (colored dots and squares) are in good agreement with model results (black dots and squares) and can be fitted by a sharp Hill function (Hill coefficient = 6.09). For each cell type, inhibition of the ephrin pathway by the injection of Eph3ΔC shifts the representative points to the right. While the effect of this shift on *Otx* expression is negligible for the a6.5, a6.6 and a6.8 cell types that are quite far from the threshold, it moves the a6.7 from the steep part of the curve to the plateau. Thus, *Otx* expressions in the a6.7 and a6.5 cell types become similar.

The highly nonlinear relation between *Otx* expression and cell surface contacts in 32-cell stage *Ciona* embryos does not rely on an ultrasensitive ERK signaling, but rather on a highly nonlinear relation between ERK activity and expression of the *Otx* gene. This threshold-like behavior is made possible by the presence of an inhibitor of *Otx* expression, ERF2 (Williaume et al., 2021). The key role of this transcriptional inhibitor is visualized in Fig. 3F. Using the values of parameters calibrated on experimental data (Table 1), the levels of active, phosphorylated ETS1/2 activator (green curve in Fig. S2B) and of active, unphosphorylated ERF2 inhibitor (red line) are plotted as a function of Erk*. These curves are sharp (n_H_=6.3 for the two curves) despite the Michaelis-Menten kinetic functions, because the phosphorylation and dephosphorylation reactions of the two transcription factors are not far from saturation (K_MM1_=K_MM2_= K_MM3_=K_MM4_=0.05), a situation close to zero-order ultrasensitivity (Goldbeter and Koshland, 1981). As a consequence of the combination of these two relatively sharp functions, the relation between *Otx* expression and ERK activity is characterized by a Hill coefficient of 6.3 (Fig. 3F). Interestingly, when the effect of the inhibitor is not considered in the model (I = 0), the curve is not only shifted to the left because of the smaller effective binding constant of the activator, but also becomes much smoother (Hill coefficient = 2.6). This theoretical result is in line with the overall increase and smoother behavior of *Otx* expression in embryos injected with an ERF2 anti-sense morpholino oligo (compare Fig. 3A and Fig. 3E). We thus conclude that the sharp relation between expression of the *Otx* gene and ERK activity is possible because the ERK pathway both promotes the expression of *Otx* and mitigates the inhibition of this expression.

### Sensitivity analysis

We next conducted a sensitivity analysis to assess the robustness of the model and to identify the parameters most affecting the model’s behavior. The influence of varying the value of each parameter by ±20% on the steady states of ERK activity (Erk*) and *Otx* expression (O) in the four cell types is presented in Fig. 4 (see also Table 1 for the definition of each parameter). It is clear that ERK activation (Fig. 4A) and *Otx* activation (Fig. 4B) are more sensitive to changes in the values of parameters affecting the SOS pathway (indicated in blue) than the p120RasGAP one (indicated in red). This extends our previous conclusion that the control exerted by the FGF-receptor pathway is stronger than that exerted by ephrin/Eph. The model is also quite sensitive to the value of *K*_*erk*_ that represents the fraction of Ras-GTP leading to half-maximal activation of the ERK pathway. Given that this parameter controls the entire ERK pathway, which is modeled by a single Hill function (Eq 1), it is intuitively logic that its value much affects the final value of Erk*. Expression of *Otx* is dramatically affected in the a6.7 cell type when parameters promoting ERK activation are modified (Fig. 4B), in agreement with the position of this cell on the (Erk*, O) curve discussed in the previous section. Globally, the model is very robust to changes in the value of parameters describing the relation between Erk* and O (Fig. 4C), except again for the a6.7 cells. For this cell type indeed, values of O are much influenced by the rates of phosphorylation (*k*_*MM3*_) and dephosphorylation (*V*_*MM4*_) of the ETS1/2 activator. It should be noted, however, that this sensitivity analysis does not explore situations in which the relation between Erk* and *Otx* expression is not ultrasensitive since 20% changes around the standard *K*_*MMi*_ values (0.05) still correspond to a zero-order regime. Most importantly, the a6.5>a6.7>a6.6>a6.8 profile of ERK activation and *Otx* expression remain in all tested parameter ranges (not shown). Thus, the mechanism of a neural induction controlled by the cell contact area is robust against changes in the values of parameters.

**Fig 4.**
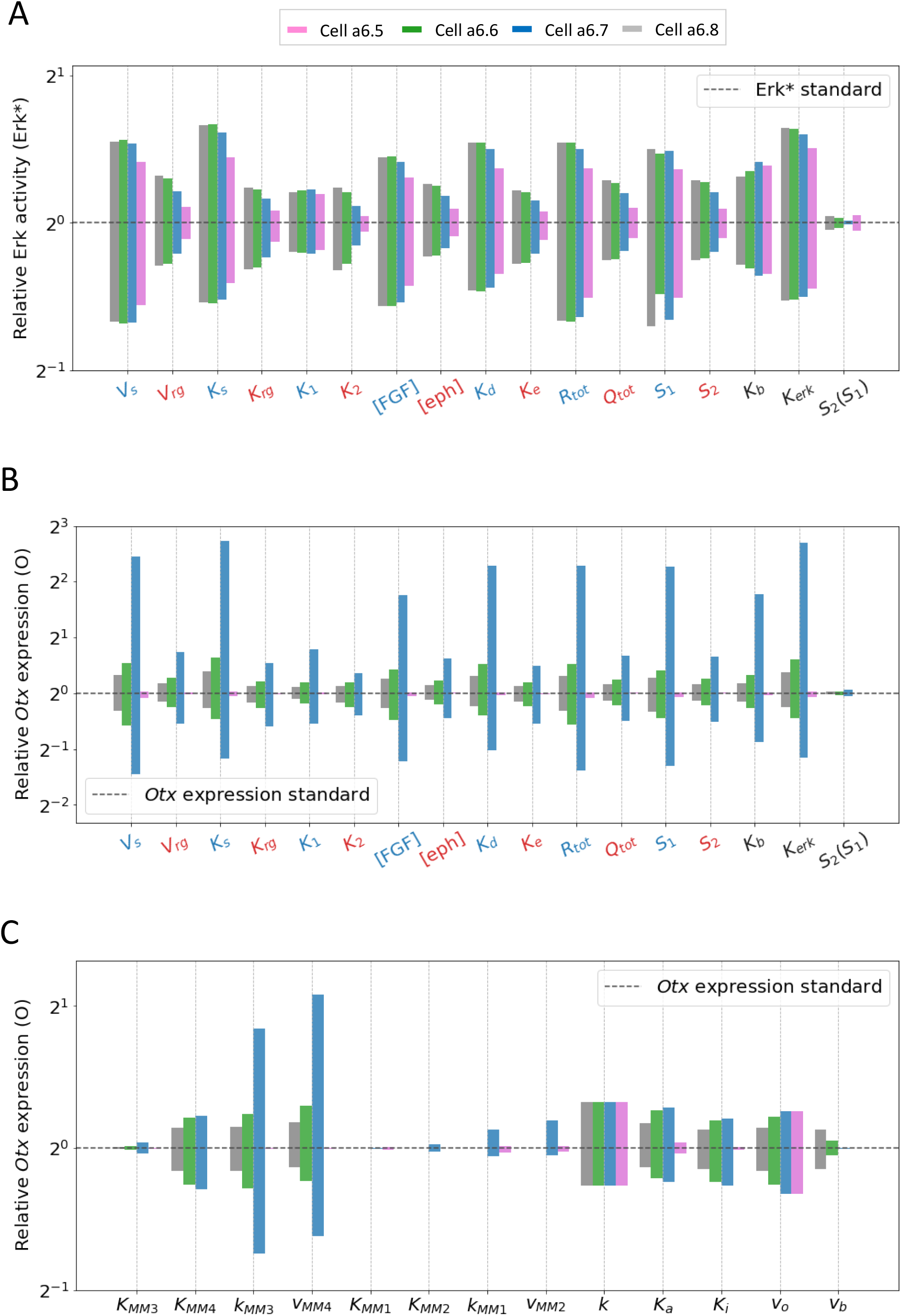
Sensitivity analysis. **(A)** Erk* values computed when varying the value of one parameter at a time by ± 20%. Shown are the ratios between the computed values and the value obtained with the default values of parameters indicated in Table 1. Parameters affecting the SOS pathway are indicated in blue, while those affecting the RasGAP one are indicated in red. Results above S_1_ and S_2_ were obtained by considering S_1_ and S_2_ as separate inputs, thus disregarding Eq 12. Results above S_2_(S_1_) were obtained by increasing/decreasing the value of the slope in Eq 12 by ± 20%, and then adjusting the intercept to best fit with experimental data. **(B)** O values computed when varying the value of one parameter at a time by ±20%. Results are obtained in the same way as for panel (A). **(C)** O values computed when varying the value of one parameter at a time by ± 20%. All panels show the ratios between the computed values and the value obtained with the default values of parameters indicated in Table 1.

### Model predictions

#### Effect of ephrin-Eph signaling

In this section, we use the model that has been calibrated on experimental data and shown to display robustness against changes in the values of parameters to make theoretical predictions. In our previous study (Williaume et al, 2021), we showed that ERK signaling is more sensitive to the cell surface contact to FGF-expressing cells (S_1_) than to FGF concentration. Quantification of this statement, in terms of differences between Hill coefficients, is shown in Fig. 5A. For each column, we use the default values of all parameters (Table 1), except the one that is indicated. In the same line, we now investigate if ERK signaling is more sensitive to cell surface contact with ephrin-expressing cells (S_2_) than to ephrin concentration itself. To this end, we computed the Hill coefficients of the Erk* vs S_2_ relation and of Erk* vs [ephrin]. As shown in Fig. 5B, ERK activity is more sensitive to cell surface contact to ephrin-expressing cells than to ephrin concentration.

**Fig 5.**
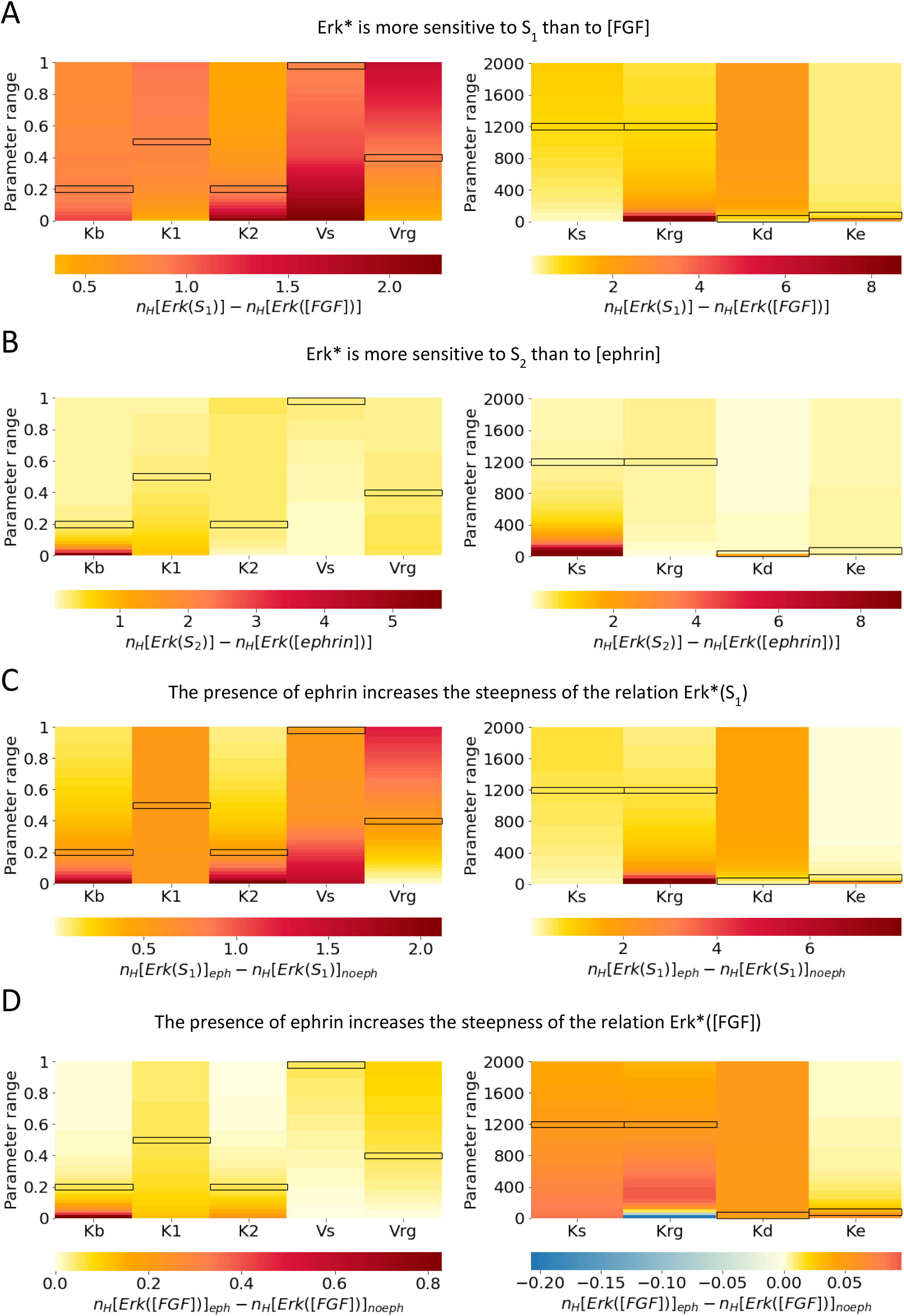
Model prediction. **(A)** Erk* values are more sensitive to S_1_ than to [FGF]. Heatmaps show the difference between the Hill coefficient of the curve Erk*(S_1_) and the Hill coefficient of the curve Erk*([FGF]). Hill coefficients obtained with default values of the parameters are 1.54 for Erk*([FGF]) and 2.39 for Erk*(S_1_). To compute Erk*([FGF]) we fixed the value of S_1_ =0.5. **(B)** Erk* values are more sensitive to S_2_ than to [ephrin]. Heatmaps show the difference between the Hill coefficient of the curve Erk*(S_2_) and the Hill coefficient of the curve Erk*([ephrin]). Hill coefficients obtained with default values of the parameters are 1.15 for Erk*([ephrin]) and 1.43 for Erk*(S_2_). To compute Erk*([ephrin]) we fixed the value of S_1_ =0.5. **(C)** The presence of ephrin increases the steepness of the relation between Erk* and S_1_. Heatmaps show the difference between the Hill coefficient of the curve Erk*(S_1_) computed in the presence of ephrin and the Hill coefficient of the curve Erk*(S_1_) computed in the absence of ephrin ([ephrin]=0.001). The Hill coefficient obtained with standard values of the parameters for Erk*(S_1_) in the absence of ephrin is 1.85. **(D)** The presence of ephrin increases the steepness of the relation between Erk* and [FGF]. Heatmaps show the difference between the Hill coefficient of the curve Erk*([FGF]) computed in the presence of ephrin and the Hill coefficient of the curve Erk*([FGF]) computed in the absence of ephrin ([ephrin]=0.001). The Hill coefficient obtained with standard values of the parameters for Erk*([FGF]) in the absence of ephrin is 1.48. To compute Erk*([FGF]) we fixed the value of S_1_ =0.5. For all panels, the Hill coefficients were computed varying the values of the parameters from 0 to 1 (K_b_, K_1_, K_2_, V_s_, V_rg_; left panels) or from 0 to 2000 (K_s_, K_rg_, K_d_, K_e_; right panels). Values of the Hill coefficients were obtained by curve fitting. The standard values of the parameters (Table 1) are highlighted with black boxes.

The model can also be used to gain a deeper understanding of the role of ephrin, which was shown to prevent ectopic expression of *Otx* in the a6.7 cell by decreasing ERK activity below the threshold level in this cell type. Simulations reveal that the presence of ephrin also increases the steepness of the relation between Erk* and the cell surface contact to FGF-expressing cells, S_1_ (Fig. 5C). Interestingly, the steepness of the relation between Erk* and [FGF] is also increased by the presence of ephrin, although this effect is less pronounced (Fig. 5D). In the two situations, the increase in steepness of the Erk* ([FGF]) (or Erk*(S_1_)) relation due to the presence of ephrin is maximal in the absence of ephrin-independent RasGAP activity (K_b_=0). The increased steepness of the Erk*([FGF]) relation is however not observed at low values of K_rg_, which defines the sensitivity of p120RasGAP to ephrin-bound receptors (Fig. 5D right). For small values of K_rg_, p120RasGAP activation is actually insensitive to ephrin, which makes the computation physiologically meaningless. Results thus show that ephrin signaling reinforces the sensitivity of the cells to changes in cell surface contact with FGF-emitting cells.

#### Comparison between FGF>FGFR >SOS and ephrin>Eph>RasGAP signaling

As qualitatively discussed previously (Williaume et al., 2021), the model indicates (Fig. 6A) that ERK signaling is more sensitive to changes in the cell surface contact to FGF-expressing cells (S_1_) than to the cell surface contact to ephrin-expressing cells (S_2_) when considering these two surfaces as independent parameters. Simulations shown in Fig. 6B indicate that, in line with this property, Erk* is more sensitive to [FGF] than to [ephrin]. It is thus clear that there is an asymmetry in the respective influences of the FGF>FGFR>SOS and of the ephrin>Eph>RasGAP pathways in controlling the possible outcome of the neural fate. Simply stated, the upregulation of one of the pathways is not equivalent to the downregulation of the other. Interestingly, even in the absence of the ephrin-independent RasGAP activity (K_b_), the Erk* vs [FGF] curve is slightly steeper than the Erk* vs [ephrin]. Indeed, Fig. 6C indicates Hill coefficients of 2.14 and 1.81 respectively for the Erk* vs [FGF] and Erk* vs [ephrin] relations. For these curves, the values of all parameters have been taken the same for the FGF and the ephrin pathway, with K_b_∼0 and S_1_ and S_2_ are both equal to 0.5. Even in these completely symmetrical conditions, the domain of values covered by the fraction of Ras-GTP (T) differs when varying FGF or ephrin concentrations (Fig. S3). In consequence, Erk* can never reach zero when increasing [ephrin] and can never reach 1 when increasing [FGF], which slightly modifies the steepness of the curves (Fig. 6C).

**Fig 6.**
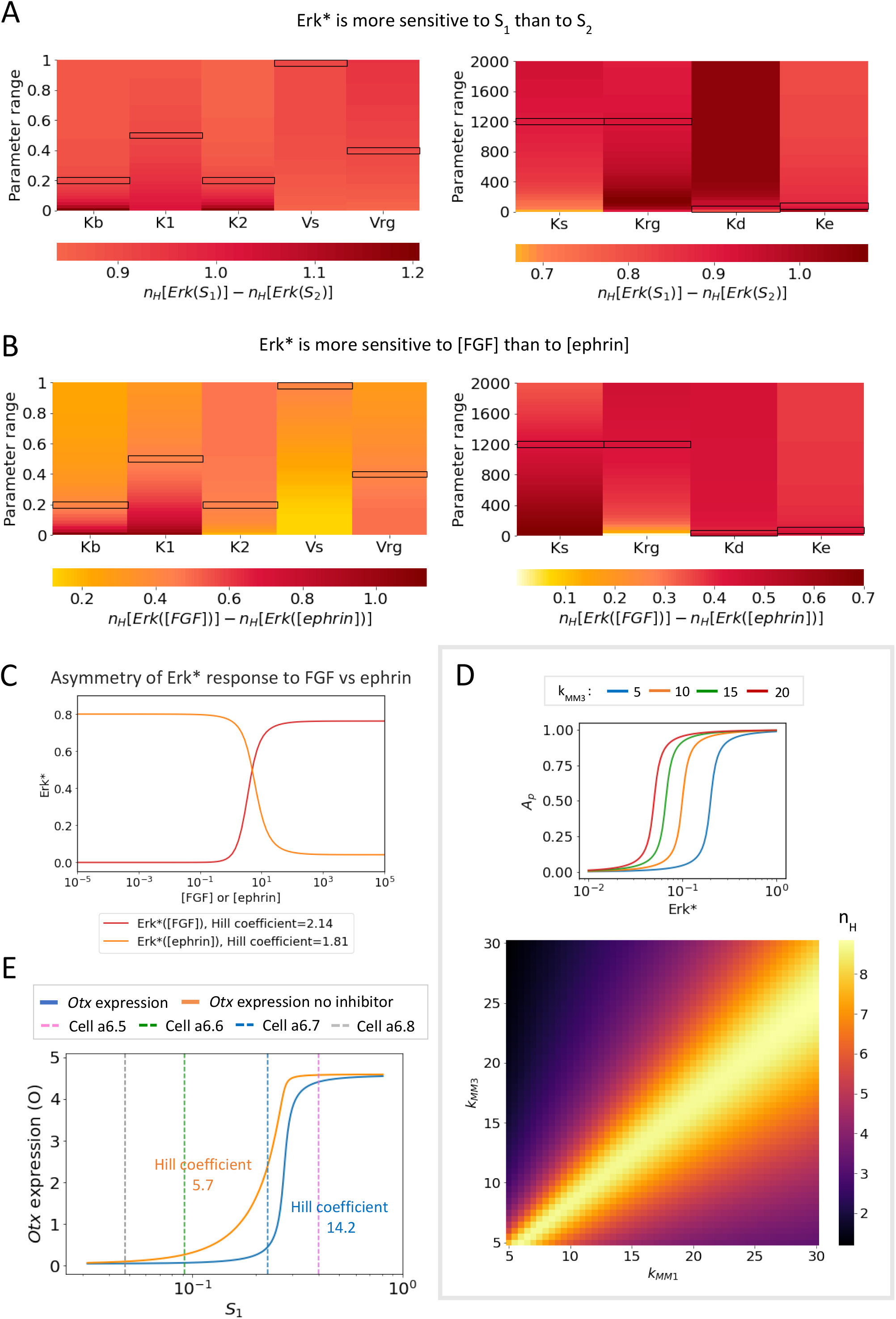
Model prediction. **(A)** Erk* values are more sensitive to S_1_ than to S_2_. Heatmaps show the difference between the Hill coefficient of the curve Erk*(S_1_) and the Hill coefficient of the curve Erk*(S_2_). Erk*(S_1_) was obtained by fixing the value of S_2_ = S* (S* =0.2) and letting S_1_ vary between 0 and 1-S*. The same procedure was used to compute Erk*(S_2_). The Hill coefficients obtained with standard values of the parameters are 1.96 and 1.05 for Erk*(S_1_) and Erk*(S_2_), respectively. **(B)** Erk* values are more sensitive to [FGF] than to [ephrin]. Heatmaps show the difference between the Hill coefficient of the curve Erk*([FGF]) and the Hill coefficient of the curve Erk*([ephrin]). To compute Erk*([FGF]) and Erk*([ephrin]) we fixed the value of S_1_ =0.5. The standard values of the parameters are highlighted with black boxes. **(C)** ERK activity (Erk*) is shown as a function of FGF concentration (in red) or ephrin concentration (in orange). Values of the parameters used: V_s_= V_rg_=1, K_1_= K_2_=0.5, K_rg_= K_s_=1200, [FGF] or [ephrin]=5, K_d_= K_e_=25, R_T_= Q_T_=2000, S_1_= S_2_= 0.5, K_b_=10^−6^. Values of the Hill coefficients were obtained by curve fitting. **(D)** Upper panel: Effect of changing the k_MM3_ in Eqs (16-17) on the relation between the fraction of phosphorylated activator, A_p_ and active ERK, Erk*. Lower panel: Heatmaps showing the Hill coefficients of the relation between *Otx* and Erk* when changing the values of the k_MMi_ in Eqs (16-17). **(E)** Relation between the level of *Otx* expression predicted by the model (O) and cell surface in contact with FGF-expressing mesendoderm cells (S_1_) in the presence (blue curve, Hill coefficient = 14.2) or in the absence (orange curve, Hill coefficient = 5.7) of the inhibitor I. Dashed vertical lines represents the mean values of S_1_ for each cell type. The Hill coefficients were computed using relation (23).

#### Effects of changing the kinetics of regulation of Otx expression

Finally, we explored the influence of the kinetics of the regulators of *Otx* expression on its switch-like behavior. It is highly sensitive to the steepness of the phosphorylation/dephosphorylation cycle of the activator and inhibitor, measured by the values of the K_MMi_ (Fig. S4). Small values of these Michaelis-Menten constants are needed to get large Hill coefficients in the relation between *Otx* expression and ERK activity. These small values allow for zero-order ultrasensitivity (Goldbeter and Koshland, 1981). Cooperativities in the effects of the activators and inhibitors on *Otx* expression could play the same role but have not been considered in the model. The effects of the two regulators are however not symmetrical because the switch-like behavior cannot be obtained if only the inhibitor phosphorylation cycle has a steep dependence on ERK activity (low values of K_MM1_ and K_MM2_). Concerning the rates of phosphorylation, the switch-like behavior is optimized when rates are similar for the activator and the inhibitor (lower panel of Fig. 6D). In this case, A_p_ and I would be sensitive to similar ranges of Erk* values (see upper panel of Fig. 6D for A_p_). The relation between *Otx* expression (O) and the cell surface in contact with FGF-expressing mesendoderm cells (S_1_) computed by the model is shown in Fig. 6E. The Hill coefficients in the presence (14.2) and absence (5.7) of inhibitor are about two times larger than those characterizing the relation between O and Erk* (6.3 and 2.6, respectively, see Fig. 3F). Thus, the combination between a moderately nonlinear relation between Erk* and S_1_ (Fig. 2B) and a steep dependence of O on Erk*, favored by the presence of the ERF2 inhibitor, allows for a quasi all-or-none dependence of the level of *Otx* expression on the cell surface contact. The relation between *Otx* expression and S_1_, which would be difficult to obtain experimentally, also helps visualize how small variations in the values of the cell surface contact of a6.7 cells, which most probably occur in some embryos, would allow them to occasionally express *Otx*, in agreement with the experimental observations (Ohta and Satou, 2013). As well as slight changes in cell surface contact, it is also possible that cell-cell heterogeneity in parameter values from one embryo to another may contribute to the increased variability in *Otx* expression levels observed in in a6.7 cells (Figure 4B). In short, the position of the a6.7 cell on the slope of the response curves make it more prone to variation in *Otx* expression compared to the other cells.

## Discussion

Modelling is much used in developmental biology to unravel the molecular mechanisms driving cell specification (see, for example: Rué and Garcia-Ojalvo, 2013; Csikasz-Nagy et al., 2022). The model developed in this study supports the previously proposed hypothesis that the area of cell surface contacts are key determinants for ascidian embryogenesis (Guignard et al., 2020). This situation contrasts with the role of ligand concentration gradients which are widely employed in other model systems (Sagner and Briscoe, 2017). In the specific case of ascidian neural induction, a full mathematical description of the molecular mechanism, from activation of FGF and ephrin receptors to *Otx* expression, leads to a good quantitative agreement with *in vivo* observations, both in normal and perturbed conditions. These results, which display strong robustness with respect to the values of parameters, confirm that enough information is encoded in the area of cell surface contact with two signaling molecules to determine the epidermis *versus* neural fate induction in the ascidian 32-cell stage embryo. The model moreover reveals that the two signals do not play symmetrical roles. While FGFR act as the primary determinant of ERK activation in each cell, differential ephrin/Eph receptor activation increases the sensitivity to FGF signaling. The somewhat secondary role of ephrin/Eph signaling is due to the existence of an ephrin-independent RasGAP activity. However, even in the absence of ephrin-independent RasGAP activity, there is an intrinsic asymmetry in the respective impacts of the Ras-GTP producing and reducing enzymes, as illustrated in Fig. 6C. Finally, the larger role played by FGFR than by ephrin/Eph receptors in the control of ERK activity is also due to the small Ras-GTP/Ras-GDP ratio that is predicted by the model. As a consequence, SOS possesses a larger amount of available substrate, which makes changes in its level of activity more impactful on ERK activation levels than p120RasGAPs. As well as increasing the sensitivity of the response to FGF signals, ephrin/Eph signals also dampen ERK activation levels in all cells. This is critical in a6.7 cells in which ERK activation level is near the threshold for *Otx* expression.

The best agreement between modeling and experimental results is obtained for values of parameters leading to a predicted modest level of ERK activation in ectoderm cells. In particular, modeled ERK activation level in the a6.5 cell under normal conditions is assumed to be lower than 25% of its maximum activation level (Figure S1A). This prediction is in line with the observation that nuclear dpERK IF signal levels detected in a6.5 cells can increase by a factor of ∼2 in a6.5 cells of ephrin-inhibited embryos or by a factor of ∼5 in maximally responding cells of ectodermal explants treated with high concentrations of exogenous FGF (Williaume et al., 2021). Thus, experimental observations support the model prediction that levels of ERK activation in individual cells is far from maximum.

In the model, sub-maximal ERK activation originates from the hypothesis that FGF concentration in the 32-cell stage ascidian embryo is smaller than the K_D_ of FGF binding to its receptor. This holds for all ectoderm cells, because the FGF concentration is assumed to be the same for each cell. Differential levels of signaling between the cells is, in contrast, due to different areas of cell surface in contact with FGF-secreting mesendoderm cells. Assuming a homogenous distribution of FGF receptors, signaling regulated by the number of receptors appears to be more sensitive than signaling based on the modulation of ligand concentration (Fig. 5A; Williaume et al., 2021). That the number of possibly activatable receptors, rather than the agonist concentration, plays a primary role in the signaling pathway has been observed in other instances. For example, the existence and frequency of Ca^2+^ oscillations in cells expressing the glutamate metabotropic receptor of type 5 (mGluR5) is controlled by the density of these receptors in the plasma membrane of stimulated cells rather than by the glutamate concentration (Dupont et al., 2011). In addition, during cellularization in early *Drosophila* embryos the activation of ERK in the ventral ectoderm in response to the EGFR ligand, Spitz, can also act as a sensor of the numbers of receptors available since halving the number of EGFRs results in reduction of ERK activation levels by approximately half (Lim et al., 2015). In the invariantly cleaving early ascidian embryo, control by the surface-area determined number of receptors was predicted, in conjunction with binary induction outputs, to be sufficient to control specification in early embryos without additional layers of regulation (Guignard et al, 2020).

Some assumptions of the model still require further investigation. The dispersion of the nuclear dpERK IF signals and *Otx* smFISH spots is much larger in the experiments than those predicted by the model. In the model, the dispersion of the points is assumed to have two qualitatively different origins. First, for each cell type, ERK signaling slightly differs from one cell to the other because cells slightly differ by their cell surface area in contact with FGF-expressing cells (S_1_) and ephrin-expressing cells (S_2_) (Fig. 1C and 1D). Second, to compare the normalized values of ERK activation levels computed with the model to experimentally obtained nuclear dpERK IF signals, we took into account the background level of signal. The variation on this signal, which we identified to the variation in ERK activation levels in the less responding a6.8 cell, must thus also be considered (see the section “Estimation of the uncertainties” below). This explains why the error bars of the simulated values of the nuclear dpERK IF signals and nuclear *Otx* smFISH spots exceed the dispersion of the individual points. However, in experimental measurements, if we consider a6.8 levels of ERK activation as background and normalize dpERK IF measurements in the remaining cells of each embryo half to the same side a6.8 value, the large variation in points persists (not shown). Thus, the variation in our experimental values does not only result from differential basal levels of fluorescence. The larger dispersion of experimentally measured values suggests the existence of cell-to-cell differences, other than cell surface areas, that are not considered in the model. In the same line, it is interesting to note that when fitting model outputs to experiments following inhibition of ERF2 function, a larger than usual background subtraction is required, indicating a larger than predicted basal level of *Otx* expression, even in cells with low ERK activity, when repression via ERF2 is lifted. Activation of *Otx* therefore likely involves factors in addition to ERF2/ETS1/2, with Gata.a being the prime candidate currently under investigation. The presence of additional factors is not considered in the model.

Another assumption of the model relates to the origin of the ultrasensitive relation between ERK and *Otx* expression. While we have shown that the presence of the ERF2 inhibitor further increases the steepness of the relation, the already non-linear relation between ERK and *Otx* expression (n_H_= 2.6, see Fig. 3F) was postulated to arise from the presence of zero-order ultrasensitivity in the relationship between Ets1/2 and ERK* (Goldbeter and Koshland, 1981). There is actually no experimental evidence for such phenomenon in the regulation of *Otx* expression, although it has been reported to be the case for other developmental-related processes (Melen et al., 2005). Other possibilities, such as cooperative binding of the effectors, could be involved and play the same role as ultrasensitivity from a mechanistic point of view, without changing the general conclusions drawn from the present study. Further investigation of the mechanism by which a graded ERK signal is converted into a bimodal transcriptional output of immediate-early genes during ascidian neural induction would improve our general knowledge of the input-output relationship of the numerous ERK-activated processes.

## Model

We developed a computational model to describe the regulation of *Otx* expression by the FGF- and ephrin-regulated ERK pathway during ascidian neural induction. The model results from the combination and parametrization of the three independent modules that were developed in our previous work to help understand how signaling inputs are integrated to activate ERK signaling on one hand, and how ERK signaling modulates *Otx* expression on the other hand (Williaume et al., 2021). Here we combine these modules and calibrate the values of the parameters to compare the outcome of the full model with experimental results.

### Model of ERK activity

We assumed that FGF and ephrin act as short-range ligands, with ephrins being membrane-tethered ligands, and that the concentration of extracellular FGF ([FGF]) between mesendoderm and ectoderm cells is uniform. Similarly, we assumed the concentration of extracellular ephrin ([ephrin]) to be constant. In line with experimental observations (Williaume et al., 2021), we considered that the density of both FGF and Eph receptors on the plasma membrane of ectoderm cells is uniform. Thus, the number of receptors in contact with the ligand is different for each ectoderm cell, since it is proportional to the area of cell surface in contact with FGF- or ephrin-expressing cells.

The activation (phosphorylation) of the extracellular signal-regulated kinase, ERK, is induced by Ras-GTP (see Fig 1A) via a cascade of phosphorylations that is not considered explicitly. Assuming that the total amount of Ras-GDP and Ras-GTP is conserved, we described the relation between the fraction of active, doubly phosphorylated ERK (*Erk*^∗^) and the fraction of Ras bound to GTP (*T*) as a Hill function, i.e.:

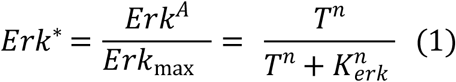

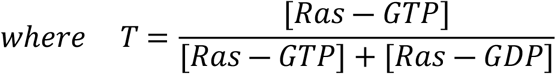

*Erk*_*max*_is the maximal level of ERK activity (*Erk*^*A*^), *n* is the Hill coefficient and *K*_*erk*_ is the fraction of Ras-GTP leading to half maximal ERK activity.

The evolution equation for the fraction of Ras-GTP is (see Fig.1A):

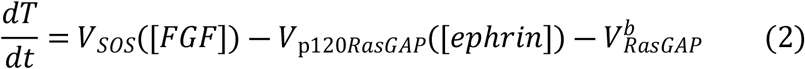

where *V*_*SOS*_([*FGF*]) is the conversion rate from Ras-GDP to Ras-GTP (mediated by the FGF dependent SOS activity), *V*_p120*RasGAP*_([*ephrin*]) is the conversion rate from Ras-GTP to Ras-GDP (mediated by the ephrin dependent p120RasGAP activity) and 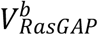 is the conversion rate from Ras-GTP to Ras-GDP mediated by ephrin-independent RasGAP activity. We will now describe these three processes.

#### SOS-mediated conversion from Ras-GDP to Ras-GTP

Ras-GDP is transformed into Ras-GTP by SOS following Michaelian kinetics. Thus, the conversion rate *V*_*SOS*_ can be written as:

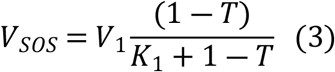

where *V*_1_ is the normalized maximal rate of conversion of Ras-GDP to Ras-GTP by SOS and *K*_1_ is the normalized half-saturation constants of SOS for its substrate Ras-GDP. To take the cell-contact dependent number of ligand-bound FGF receptors into account, we considered that SOS must be in its active state SOS* (namely membrane-recruited) to promote the formation of Ras-GTP.

The reaction for the activation of SOS depends on the number of FGF-bound receptors, *R*_*b*_. So, we re-write the rate *V*_1_ as:

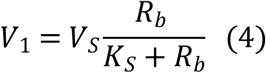

where *V*_*S*_ is the maximal rate of SOS activation and *K*_*S*_ is the number of FGF-bound receptors leading to half of the maximal SOS activity. We do not model explicitly the heparan-sulfate dependent dimerization of bound FGF receptors that precedes activation of SOS. *R*_*b*_ is given by:

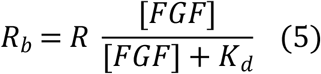

where [*FGF*] is the extracellular concentration of FGF, *R* is the number of receptors possibly in contact with FGF, and *K*_*d*_ is the binding constant of FGF to its receptor. Since we assumed that the density of FGF receptors on the ectoderm cell membrane is constant, the number of receptors in contact with FGF can be written as:

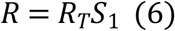

where *R*_*T*_ is the total number of FGF receptors present on a cell, and *S*_1_ is the fraction of cell surface exposed to FGF.

#### Ephrin dependent RasGAP-mediated conversion from Ras-GTP to Ras-GDP

The conversion of Ras-GTP to Ras-GDP is mediated by p120RasGAP following Michaelian kinetics. Thus, the conversion rate *V*_*Ras*―*GAP*_ can be written as:

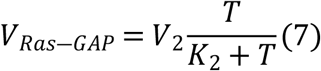

where *V*_2_ is the normalized maximal rate of conversion of Ras-GTP to Ras-GDP by p120RasGAP and *K*_2_ is the normalized half-saturation constant of p120RasGAP for its substrate Ras-GTP. To take the cell-dependent number of Eph receptors into account, we considered that p120RasGAP must be in an active state p120RasGAP* (membrane-recruited) in order to promote the formation of Ras-GDP.

The activation of p120RasGAP depends on the number of receptors bound to ephrin (*Q*_*b*_). Thus, we re-write the transformation rate *V*_2_ as:

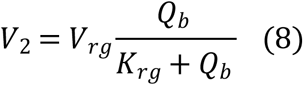

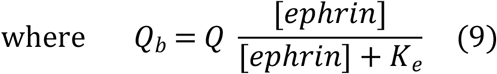

*V*_*rg*_ is the maximal rate of p120RasGAP activation, *K*_*rg*_ is the number of ephrin-bound Eph receptors leading to half of the maximal p120RasGAP activity. *Q* is the number of Eph receptors in contact with ephrin, [*ephrin*] is the ephrin concentration and *K*_*e*_ is the binding constant of ephrin to its receptor.

Since we assumed that the density of Eph receptors on the ectoderm cell membrane is constant, the number of receptors in contact with ephrin can be written as:

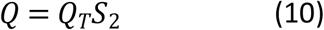

where *Q*_*T*_ is the total number of Eph receptors present on a cell, and *S*_2_ is the fraction of cell surface exposed to ephrin.

#### Ephrin independent RasGAP-mediated conversion from Ras-GTP to Ras-GDP

The presence of basal GAP activity was considered since transcriptome datasets (Brozovic et al., 2018) indicate the presence of at least three further RasGAPs in the early ascidian embryo (*IQGAP1/2/3, Neurofibromin*, and *RASA2/3*). The conversion from Ras-GTP into Ras-GDP due to contact surface independent RasGAP activity is modeled by a linear reaction rate *K*_*b*_. Thus, the conversion rate 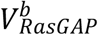 can be written as:

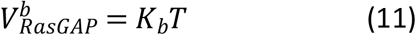

#### Empirical relation between fractions of cell surfaces in contact with FGF and with ephrin

To account for the fact that the number of Eph receptors exposed to ephrin changes together with the number of FGF receptors exposed to FGF, i.e., that cells that have the highest number of FGF receptors in contact with FGF (R), have also the lowest number of Eph receptors in contact with ephrin (*Q*), we used the experimental data from Fig. 1D. Plotting *S*_2_ as a function of *S*_1_ and fitting the experimental data with a linear function, we obtained the following relation between *S*_1_ and *S*_2_:

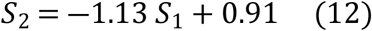

Everything considered, the evolution equation for *T* (Eq 2) becomes:

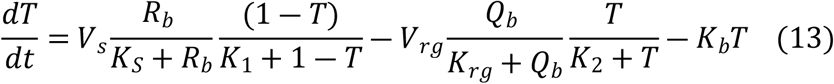

We solved the equation for *T* at equilibrium, to get *T* as a function of [*FGF*] or as a function of *S*_1_. We then computed the fraction of active ERK using Eq 1.

Usually, the quantity measured experimentally is the level of dpERK fluorescence. Thus, to compare the simulations results with experiments we considered that the level of ERK fluorescence (*Erk*^*f*^) is a linear function of ERK activity (*Erk*^∗^):

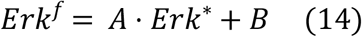

Where *A* is the maximal ERK fluorescence and *B* is the basal level of dpERK fluorescence. Taking into account that Erk* computed with the model was close to zero in the a6.8 cell type, we chose B as the average level of fluorescence in the a6.8 cell type in each experiment. A was defined empirically in order to obtain the best fit to the data.

### Model of Otx expression

*Otx* expression is regulated by dpERK (Fig. 1A). A minimal enhancer of the *Otx* gene, named the *Otx* a-element, contains 3 GATA and 2 ETS (E26 transformation-specific)-binding sites required for neural-specific expression in the ectoderm lineages via Gata.a and ETS1/2 transcription factors (Bertrand et al., 2003; Rothbacher et al., 2007).

The Ets2 repressor factor 2 (ERF2) (Yagi et al., 2003) recognizes the same binding site as the transcriptional activator ETS1/2 and its activity is negatively controlled by ERK via nuclear export (Le Gallic et al., 1999; 2004; and Weaver et al, 2022). Thus, we considered that *Otx* expression is enhanced by the stimulation of an activator (A; e.g. ETS1/2) and is inhibited by the repression of an inhibitor (I; e.g. ERF2). The phosphorylation of both the activator and the inhibitor is controlled by ERK. Thus, we modeled *Otx* expression with the following evolution equation:

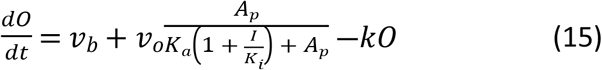

where *v*_*b*_ is the basal *Otx* expression rate, *v*_*b*_ + *v*_*o*_ is the maximal *Otx* expression rate and *k* is the degradation rate for *Otx. K*_*a*_ is the half saturation constant for the activator *A*_*p*_ while *K*_*i*_ is the dissociation constant for the inhibitor. In Eq 15, it is considered that ERF2 (*I*) is a competitive inhibitor of ETS1/2 (*A*). Thus, there is an effective half-saturation constant that depends on the concentration of *I*, which is equal to 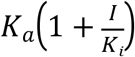. When *I* is large, this effective constant is much larger than *K*_*a*_ and, when *I* is small, this effective constant approaches *K*_*a*_. The evolution equations describing the phosphorylation and dephosphorylation of ETS1/2 and ERF2 are:

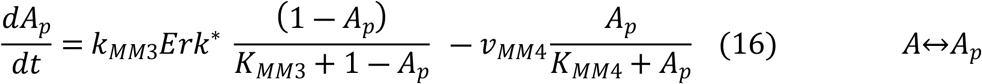

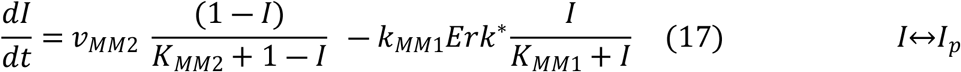

where *A*_*p*_ represents the fraction of active, phosphorylated activator; (1 ― *A*_*p*_) the fraction of inactive, unphosphorylated activator; I the fraction of active, unphosphorylated inhibitor and (1 ― *I*) the fraction of inactive, phosphorylated inhibitor. For the two phosphorylation reactions, the rate constants *k*_*MM*3_ and *k*_*MM*1_ of *A* and *I* phosphorylation are multiplied by the fraction of doubly phosphorylated ERK (Erk*). *v*_*MM*4_ and *v*_*MM*2_ are the maximal rates at which *A*_*p*_ and *I*_*p*_ are dephosphorylated. *K*_*MM*3_ and *K*_*MM*1_ are Michaelis-Menten constants for the phosphorylation of *A* and *I*, respectively, and *K*_*MM*2_ and *K*_*MM*4_ are Michaelis-Menten constants for the dephosphorylation of *I*_*p*_ and *A*_*p*_, respectively. *K*_*MM*3_ and *K*_*MM*4_are normalized with respect to the total activator concentration, while *K*_*MM*2_ and *K*_*MM*1_ are normalized with respect to the total inhibitor concentration.

Since the quantity measured experimentally is the counts of smFISH *Otx* spots, to compare the simulations results with experiments we considered that the level of counts of smFISH *Otx* spots (*Otx*_*smFISH*_) is a linear function of the level of *Otx* expression (*O*):

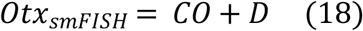

where *C* is the maximal number of smFISH *Otx* spots and *D* is the basal smFISH *Otx* spot count. Taking into account that the value of *O* computed with the model was close to zero in the a6.8 cell type, we chose *D* as the *Otx* smFISH count in the a6.8 cell type in each experiment. C was defined empirically in order to obtain the best fit to the data.

To obtain an expression for *Otx* expression (O and Otx_smFISH_) as a function of the inputs of the signaling cascade ([*FGF*] and *S*_1_) we combined the models of ERK activity and of *Otx* expression presented in the previous sections. We computed ERK activity using Eq 1 and we used this as the input (Erk*) to compute the fraction of active ETS1/2 (*A*_*p*_) and active ERF2 (*I*) using Eq 16 and 17. Then we computed the level of *Otx* expression by solving Eq 18. All equations are solved at steady state.

Equations are solved analytically at steady-state and solutions evaluated for all investigated sets of values of parameters with codes written in Python. Codes are available at https://github.com/rossanabettoni/Model-of-neural-induction-in-the-ascidian-embryo

### Experimental methods

All experimental results are taken from our previous work (Williaume et al., 2021).

### Estimation of uncertainties ERK fluorescence

Given a complete set of parameter values including S_1_ and S_2_, there is no uncertainty on the deterministically modeled value of ERK activity (*Erk*^∗^). However, to compare with experimental data, we computed the average of 25 values of *Erk*^∗^ (*Erk*_*M*_), corresponding to 25 couples of (S_1_, S_2_) values of individual cells. The uncertainty on this computed average activity (Δ*Erk*_*M*_) is the standard deviation.

Experimentally, ERK activation levels are quantified by immunofluorescence (IF) signals. As described above (Eq (14)), we assumed that IF signals are a linear function of *Erk*^∗^:

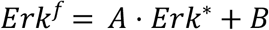

where B is the basal value of IF signals.

In the absence of other information, we postulated that the uncertainty on B (*ΔB*) is the same as the standard deviation of the experimental values of dpERK IF signals measured in the a6.8 cells. Although it may be unlikely that there is zero variability between the true value of ERK activation level and the measured fluorescence intensity in terms of slope, we have no means to measure or estimate this. Therefore, we considered that there is no uncertainty on A. Thus, the uncertainty Δ*Erk*^*f*^ is equal to Δ*B* for each cell, i.e. for each couple of (S_1_, S_2_) values. In Fig. 2A and S1C the uncertainties Δ*Erk*^*f*^ for the single cells are not shown, for clarity reasons. The average ERK fluorescence for a given cell type is computed as:

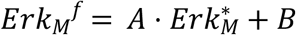

with the corresponding uncertainty:

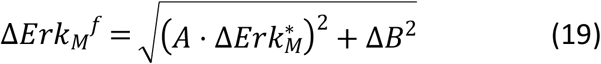

This uncertainty is indicated on the figure panels as the model error bars.

**Ratios between ERK fluorescence levels in treated and non-treated cells (for figures 2C and S1C)**

The ratio between the level of ERK fluorescence in the injected side of the embryo and the level of ERK fluorescence in the control side of the embryo (for a single cell) is:

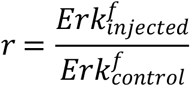

With uncertainty:

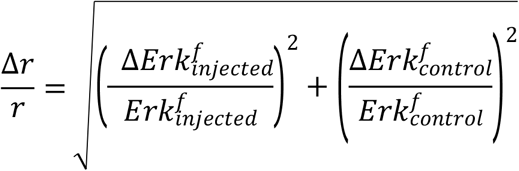

where 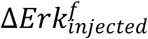 and 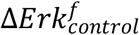 are equal to Δ*B*. In Fig. 2C and S1D the uncertainties Δ*r* for the single cells are not shown.

The average of the ratios for each cell type was computed as:

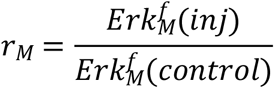

With corresponding uncertainty:

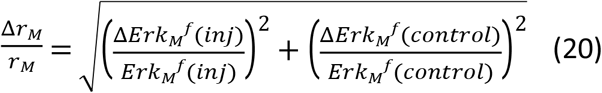

in which 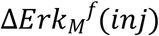and 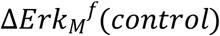 are given by Eq. (19).

This uncertainty is indicated on the figure panels as the model error bars.

### *Otx* smFISH spot counts

Given a complete set of parameter values including S_1_ and S_2_, there is no uncertainty on the deterministically modeled value of *Otx* expression (*O*). However, to compare with experimental data, we computed the average of 25 values of *O* (*O*_*M*_), corresponding to 25 couples of (S_1_, S_2_) values. The uncertainty on this computed average activity (Δ*O*_*M*_) is the standard deviation.

Experimentally, *Otx* expression is quantified by the *Otx* smFISH spot counts. As described above (Eq (18)), we assumed that fluorescence is a linear function of *O*:

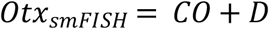

where D is the basal value of *Otx* smFISH spot counts.

In the absence of other information, we postulated that the uncertainty on D (*ΔD*) is the same as the standard deviation of the experimental values of *Otx* smFISH spot counts measured in the a6.8 cells. We considered that there is no uncertainty on C.

Thus, the uncertainty Δ*Otx*_*smFISH*_ is equal to Δ*D* for each cell, i.e. for each couple of (S_1_, S_2_) values. In the Fig. 3A, 3B, 3C and 3E the uncertainties Δ*Otx*_*smFISH*_ for the single cells are not shown, for clarity reasons.

The average *Otx* smFISH spot counts for a given cell type is computed as:

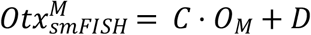

with the corresponding uncertainty:

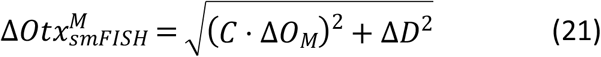

This uncertainty is indicated on the figure panels as the model error bars.

### Estimation of the Hill coefficients

When not mentioned explicitly, Hill coefficients were obtained by fitting the curves with Hill functions.

Since the fit of the curve *Otx* expression (O) as a function of Erk* in the absence of the inhibitor (Fig. 3F) was poor, we used the fact that Hill coefficients can be calculated in terms of potency as:

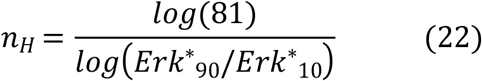

where Erk*_90_ and Erk*_10_ are the levels of ERK activity needed to produce 90% and 10% of the maximal *Otx* expression, respectively.

Similarly, we computed the Hill coefficient between *Otx* expression (O) and S_1_ in Fig. 6E as:

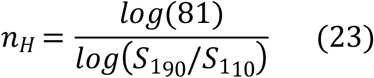

where 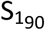 and 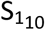 are the fractions of cell surface in contact with FGF needed to produce 90% and 10% of the maximal *Otx* expression, respectively.

## Acknowledgments

RB is supported by a FRIA fellowship. GD is Research Director at the Belgian “Fonds National pour la Recherche Scientifique” (FRS-FNRS) and acknowledges financial support from the ARC project “Noise sensitivity of gene regulatory networks underlying cell fate specification” financed by the Université libre de Bruxelles (ULB). The team of HY is supported by the Centre National de la Recherche Scientifique (CNRS), Sorbonne University, the Fondation ARC pour la Recherche sur le Cancer (PJA 20131200223) and the Agence Nationale de la Recherche (ANR-17-CE13-0003-01).

## Notes

### Competing Interest Statement

The authors have declared no competing interest.

